# Neural similarity between overlapping events at learning differentially affects reinstatement across the cortex

**DOI:** 10.1101/2022.08.18.504184

**Authors:** Melissa Hebscher, Wilma A. Bainbridge, Joel L. Voss

## Abstract

Episodic memory often involves high overlap between the actors, locations, and objects of everyday events. Under some circumstances, it may be beneficial to distinguish, or differentiate, neural representations of similar events to avoid interference at recall. Alternatively, forming overlapping representations of similar events, or integration, may aid recall by linking shared information between memories. It is currently unclear how the brain supports these seemingly conflicting functions of differentiation and integration. We used multivoxel pattern similarity analysis (MVPA) of fMRI data and neural-network analysis of visual similarity to examine how highly overlapping naturalistic events are encoded in patterns of cortical activity, and how the degree of differentiation versus integration at encoding affects later retrieval. Participants performed an episodic memory task in which they learned and recalled naturalistic video stimuli with high feature overlap. Visually similar videos were encoded in overlapping patterns of neural activity in temporal, parietal, and occipital regions, suggesting integration. We further found that encoding processes differentially predicted later reinstatement across the cortex. In visual processing regions in occipital cortex, greater differentiation at encoding predicted later reinstatement. Higher-level sensory processing regions in temporal and parietal lobes showed the opposite pattern, whereby highly integrated stimuli showed greater reinstatement. Moreover, integration in high-level sensory processing regions during encoding predicted greater accuracy and vividness at recall. These findings provide novel evidence that encoding-related differentiation and integration processes across the cortex have divergent effects on later recall of highly similar naturalistic events.

## Introduction

Everyday memory frequently involves high overlap between the actors, locations, and objects of events. Whether such similarity between events helps or hinders memory recall may in large part be determined by how events are encoded. When we encounter two events with similar features, it might be beneficial for the brain to distinguish their representations, or differentiate them, to reduce interference. Alternatively, forming overlapping representations of similar events, or integration, may aid recall by linking new memories to old memories.

How and under which conditions does the brain supports these seemingly conflicting functions? Evidence for the beneficial effect of both integration and differentiation on memory exists. Functional magnetic resonance imaging (fMRI) studies using multivoxel pattern analysis (MVPA) have shown that integration, measured as reactivation of overlapping memories at encoding, is beneficial for later memory performance (Brunec et al., 2020; Chanales et al., 2019; Koen & Rugg, 2016; Schlichting et al., 2014; Wing et al., 2020). On the other hand, greater pattern distinctiveness in the cortex and the hippocampus has been shown to be beneficial for later memory recall, suggesting differentiation (Ezzyat et al., 2018; Favila et al., 2016; Katsumi et al., 2021).

Whether highly similar events are encoded in an integrated or differentiated fashion may also affect their later neural reinstatement, the retrieval-related reactivation of brain activity present during encoding (Danker & Anderson, 2010; Rissman & Wagner, 2012). We previously found that neural dissimilarity between events at encoding leads to increased reinstatement in the visual cortex, suggesting that differentiating visual features is beneficial for reinstatement (Hebscher et al., 2021). However, more work is needed to directly link encoding processes of integration and differentiation to neural reinstatement in different cortical regions.

Recent evidence suggests that the degree and content of overlap between events can affect whether they are integrated or differentiated. For instance, shared context between events leads to strengthened associative memory, suggesting integration, while dissimilar contexts lead to interference (Cox et al., 2021). Neural integration of events at encoding has also been shown to predict subsequent memory strength when there was low interference between events (Koen and Rugg, 2016). Another factor of likely importance is where in the brain these processes are being observed. Differentiation and integration of the same events can occur simultaneously across different hippocampal subregions, which can explain how we are able to remember both specific details of items and their relationships with each other (Dimsdale-Zucker et al., 2018; LaRocque et al., 2013). Here, we predicted that cortical regions outside the hippocampus may also differentially support these processes, with parietal and temporal cortices integrating high-level contextual and semantic features of events, and occipital regions differentiating lower-level visual features.

In the present study, we examined how highly overlapping naturalistic events are encoded and how their encoding affects later recall across the cortex. Participants performed encoding and retrieval phases of a memory task during fMRI scanning. At encoding, they viewed a series of short video clips depicting everyday activities with high feature overlap, including the same actor, similar contexts, and similar content, which mimics natural memory demands. At retrieval, which occurred after all videos had been encoded, participants used recall cues to mentally replay studied videos, and their objective and subjective memory for the videos was tested (Figure 1A).

**Figure 1.**
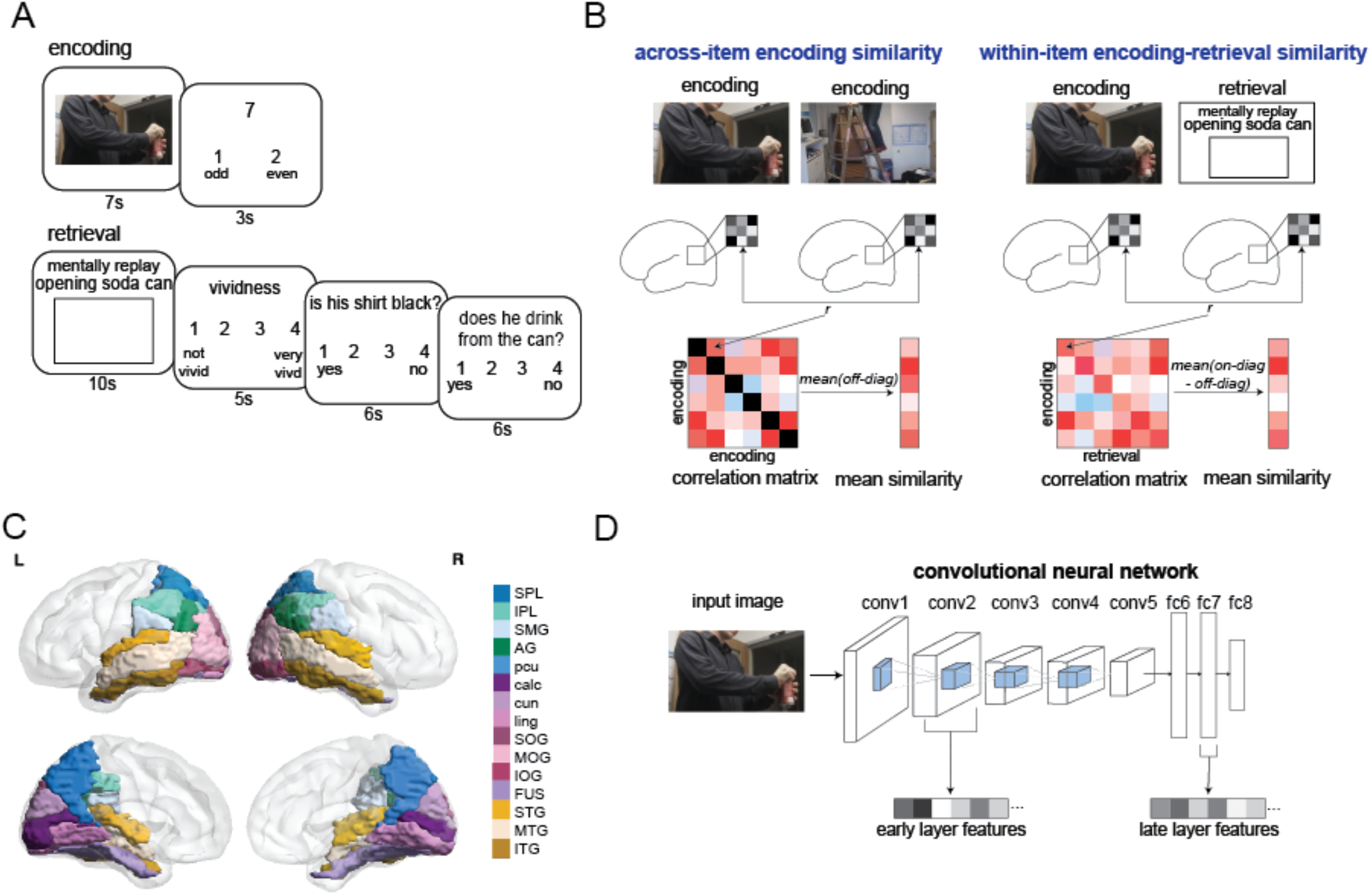
Experimental design and methods overview. (A) Example trial for the experimental task. Participants watched videos at encoding and made odd/even judgments for arbitrary numbers. At retrieval, participants were cued with titles describing each of the studied videos. Participants mentally replayed each video while viewing a blank screen and then rated the vividness of the memory and answered two true/false (yes/no) questions to test memory accuracy, including a 4-point confidence judgment. (B) Multivoxel pattern similarity analysis overview. Across-item encoding similarity (left panel) within each brain region was computed by taking the average pairwise correlation between the neural activity patterns evoked while encoding each video, reflecting average neural similarity between videos. Within-item encoding-retrieval similarity, or reinstatement (right panel) was calculated as the correlation between neural activity evoked while watching a video and while later remembering it (on-diagonal of the correlation matrix), relative to the correlation between unmatched pairs of videos at encoding and retrieval (off-diagonal). (C) ROIs shown on a glass brain. SPL=superior parietal lobule, IPL=inferior parietal lobule, SMG=supramarginal gyrus, AG=angular gyrus, pcu=precuneus, calc=calcarine gyrus, cun=cuneus, ling=lingual gyrus, SOG=superior occipital gyrus, MOG=middle occipital gyrus, IOG=inferior occipital gyrus, FUS=fusiform gyrus, STG=superior temporal gyrus, MTG=middle temporal gyrus, ITG=inferior temporal gyrus. (D) Convolutional neural network (CNN) overview. The AlexNet CNN consists of 5 convolutional layers (conv) and 3 fully connected layers (fc). One frame from the midpoint of each video was used as input for the pretrained network, and features were extracted from one early layer (conv2) and one late layer (fc7), reflecting lower- and higher-level visual features, respectively

To better understand how highly overlapping events are represented in the brain, we performed MVPA to measure similarity between videos at encoding (Figure 1B, left panel) and reinstatement of videos at retrieval (Figure 1B, right panel). We examined activity within regions of interest (ROIs) across parietal, temporal, and occipital cortices (Figure 1C), which have been implicated in the recall and reinstatement of naturalistic stimuli (Bainbridge et al., 2021; Hebscher et al., 2021; Oedekoven et al., 2017; Silson et al., 2019; Xue, 2018). To test whether the degree and content of feature overlap affects how events are encoded, we related a metric of visual similarity extracted from the AlexNet convolutional neural network (CNN) with neural activity at encoding (Figure 1D). We then examined how encoding of overlapping videos affects their later retrieval by relating neural activity at encoding activity to behavioral outcomes at retrieval and neural reinstatement of stimuli.

## Results

### Visually similar videos are integrated in cortical neural populations

To understand how the degree and content of feature overlap affects how events are encoded, we first tested whether visual similarity between videos, measured by a CNN, affects their neural similarity in the cortex, measured by representational similarity analysis (RSA) (Kriegeskorte et al., 2008). A positive relationship here would indicate that videos with high visual feature overlap are represented in overlapping patterns of neural activity, which would suggest integration. A negative relationship would indicate that videos with high feature overlap are represented in more distinct neural representations, suggesting differentiation.

Neural similarity between videos was calculated by performing a series of RSAs within cortical ROIs (Figure 1C). For each participant and each ROI, we computed the average pairwise Fisher-transformed Pearson correlation between the neural activity pattern of each video (Figure 1B, left panel). This measure, termed across-item encoding similarity, reflects the relative neural similarity of a given video to all other videos, effectively its place on a continuum from neural averageness to distinctness (See Methods). To obtain a measure of visual similarity, we extracted learned image features from the pretrained CNN AlexNet (Krizhevsky et al., 2012) (Figure 1D). We used the middle frame from each video as input to the CNN and extracted features from early (conv2) and late (fc7) layers, which are thought to correspond to visual information from early and late visual cortex, respectively (Güçlü & van Gerven, 2015; Yamins et al., 2014) (see Methods). Visual similarity was computed as the average Fisher-transformed pairwise Spearman correlation between all videos. This measure reflected the degree and content of feature overlap of stimuli, with the early layer reflecting lower-level visual feature overlap and the late layer reflecting higher-level visual feature overlap.

To assess whether visual similarity between videos affects their neural similarity, we calculated the Spearman correlation between CNN-based visual similarity and fMRI across-item encoding similarity for each participant and each ROI. Permutation tests performed at the group-level revealed that visual similarity was significantly positively associated with across-item encoding similarity in parietal, temporal, and occipital regions. These tests assessed the probability that the observed correlations are more extreme than zero, relative to a null distribution of correlation differences generated by randomly signing correlation values over 1000 permutations (See Methods). The correlation between early CNN layer visual similarity and across-item encoding similarity was significantly greater than zero in bilateral middle occipital gyrus, superior occipital gyrus, left inferior occipital gyrus, left fusiform gyrus, right inferior parietal lobe, inferior temporal gyrus, calcarine gyrus, lingual gyrus, and cuneus (all p’s < 0.05 after Bonferroni-Holm correction for multiple comparisons) (See Figure 2A) (Table 1). Late layer similarity was significantly correlated with across-item encoding similarity in right inferior temporal gyrus (p < 0.05, corrected).

**Table 1.** Visual similarity – across-item encoding similarity correlation by ROI.

| Region | Early layer (conv2) |  | Late layer (fc7) |  |
| --- | --- | --- | --- | --- |
| | $r_s$ | p-value | $r_s$ | p-value |
| L AG | 0.04 | 1.0 | 0.03 | 1.0 |
| L IPL | 0.14 | 0.076 | 0.09 | 0.1 |
| L PCU | 0.11 | 0.34 | 0.03 | 1.0 |
| L SMG | 0.10 | 0.182 | 0.1 | 0.26 |
| L SPL | 0.14 | 0.09 | 0.06 | 1.0 |
| L IOG | 0.14 | 0.03<sup>a</sup> | 0.09 | 0.058 |
| L MOG | 0.17 | 0.03<sup>a</sup> | 0.08 | 0.306 |
| L SOG | 0.16 | 0.03<sup>a</sup> | 0.08 | 0.323 |
| L calc | 0.12 | 0.09 | -0.01 | 1.0 |
| L cun | 0.04 | 1.0 | 0.01 | 1.0 |
| L fus | 0.13 | 0.03<sup>a</sup> | 0.01 | 1.0 |
| L ling | 0.09 | 0.195 | -0.04 | 1.0 |
| L ITG | 0.11 | 0.135 | 0.07 | 0.524 |
| L MTG | 0.06 | 0.566 | 0.1 | 0.058 |
| L STG | 0.05 | 0.599 | 0.04 | 1.0 |
| R AG | 0.04 | 1.0 | 0.02 | 1.0 |
| R IPL | 0.14 | 0.03<sup>a</sup> | 0.13 | 0.058 |
| R PCU | 0.09 | 0.34 | -0.0 | 1.0 |
| R SMG | 0.06 | 0.464 | 0.1 | 0.176 |
| R SPL | 0.11 | 0.195 | 0.09 | 0.252 |
| R IOG | 0.12 | 0.112 | 0.1 | 0.4 |
| R MOG | 0.15 | 0.03<sup>a</sup> | 0.11 | 0.1 |
| R SOG | 0.17 | 0.03<sup>a</sup> | 0.12 | 0.1 |
| R calc | 0.13 | 0.03<sup>a</sup> | 0.0 | 1.0 |
| R cun | 0.142 | 0.03<sup>a</sup> | 0.09 | 0.285 |
| R fus | 0.078 | 0.319 | -0.02 | 1.0 |
| R ling | 0.12 | 0.03<sup>a</sup> | -0.03 | 1.0 |
| R ITG | 0.11 | 0.03<sup>a</sup> | 0.16 | 0.03<sup>a</sup> |
| R MTG | 0.01 | 1.0 | 0.09 | 0.078 |
| R STG | -0.04 | 1.0 | 0.03 | 1.0 |

*reported p-values are Bonferroni-Holm corrected for multiple comparisons*
*$r_s$ = Spearman's rho*
*<sup>a</sup> passed Bonferroni-Holm correction for multiple comparisons*

**Figure 2.**
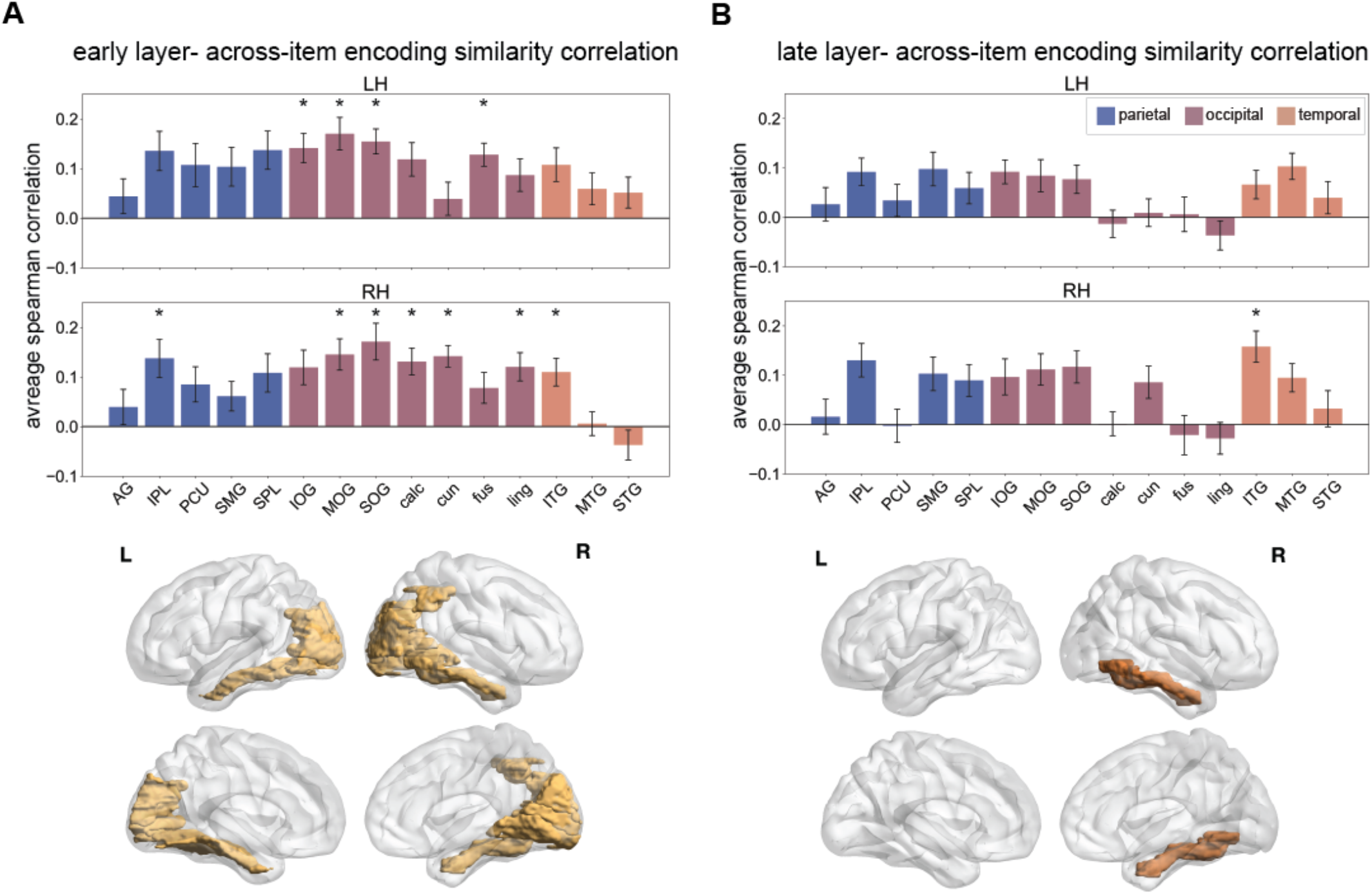
Visual similarity predicts across-item encoding similarity. A) Average within-subject correlation between visual similarity and across-item encoding similarity for an early CNN layer (conv2). Average correlations are shown for parietal, occipital, and temporal ROIs, split by hemisphere (LH – left hemisphere, RH – right hemisphere). Glass brains below show lateral (top) and medial (bottom) views of corresponding significant ROIs for left (L) and right hemispheres (R). B) Average within-subject correlation between visual similarity and across-item encoding similarity for a late CNN layer (fc7). Glass brains below show lateral (top) and medial (bottom) views of corresponding significant ROI for left and right hemispheres. Standard error of the mean plotted as error bars. Asterisks show significant regions (p < 0.05, Bonferroni corrected for multiple comparisons).

These findings suggest that similar videos are integrated with one another in the cortex, such that videos with higher visual feature overlap also show higher pattern similarity with other videos. Conversely, videos that are dissimilar tend to be differentiated, or encoded with low pattern similarity. Early layer CNN features are related to representations mainly in occipital cortex, including both lower-level (calcarine gyrus, lingual gyrus, cuneus) and higher-level visual processing regions (i.e. fusiform gyrus, occipital gyri, inferior temporal gyrus), while late layer CNN features are related to representations in inferior temporal gyrus, a high-level visual processing region.

### Across-item encoding similarity differentially predicts reinstatement

We next evaluated how encoding overlapping videos affects their later retrieval by testing whether across-item encoding similarity affects later reinstatement of videos across the cortex. We measured reinstatement of individual video clips by comparing patterns of fMRI activity between encoding and retrieval. For each participant and ROI, encoding-retrieval similarity was computed as the Pearson’s correlation of a video’s fMRI activity pattern during encoding with the corresponding activity pattern when participants attempted to mentally replay the same video during retrieval (See Figure 1B, right panel). Encoding-retrieval similarity was computed for all pairs of videos, resulting in a Fisher-transformed correlation matrix in which the diagonal reflected reinstatement of video-specific neural activity (same video at encoding and retrieval), and the off-diagonal reflected reinstatement of mismatched pairs of videos (different video at encoding and retrieval). The average correlation for the mismatched pairs (off-diagonal of the matrix) was then subtracted from the correlation for matched pairs (diagonal of the matrix). The resulting difference scores reflected the degree to which video-specific neural activity at encoding was reinstated at retrieval, relative to reinstatement of other videos. Importantly, there was no inherent perceptual similarity between encoding and recall, so reinstatement reflects memory rather than perceptual overlap. Average reinstatement scores were above zero for all ROIs other than right lingual gyrus (mean = -0.001), indicating that video-specific content was reinstated to a greater degree than content not specific to each video.

To quantify the effects of across-item encoding similarity on later reinstatement, we computed the Spearman correlation between across-item encoding similarity and reinstatement for each subject and each ROI. Permutation tests revealed this correlation was significantly different than zero for several ROIs, with the direction of this correlation differing by region. Across-item encoding similarity and reinstatement were positively correlated in temporal cortex (significant effects in bilateral inferior temporal gyrus, and right superior temporal gyrus), and negatively correlated in occipital regions (significant effects in bilateral calcarine gyrus, lingual gyrus, left middle occipital gyrus and left fusiform gyrus (all p’s < 0.05, corrected for multiple comparisons). See Figure 3, Table 2. As can be seen in Table 2, additional regions showed similar trends that did not survive correction for multiple comparisons, including positive effects in right supramarginal gyrus (p < 0.01; p = 0.06, corrected) and middle temporal gyrus (p < 0.01; p = 0.06, corrected).

**Table 2.** Across-item encoding – reinstatement similarity correlation by ROI.

| Region | $r_s$ | p-value |
| --- | --- | --- |
| L AG | 0.01 | 1.0 |
| L IPL | 0.12 | 0.364 |
| L PCU | 0.03 | 1.0 |
| L SMG | 0.12 | 0.133 |
| L SPL | -0.03 | 1.0 |
| L IOG | -0.09 | 0.432 |
| L MOG | -0.12 | 0.048<sup>a</sup> |
| L SOG | -0.11 | 0.133 |
| L calc | -0.21 | 0.03<sup>a</sup> |
| L cun | -0.07 | 0.759 |
| L fus | -0.14 | 0.048<sup>a</sup> |
| L ling | -0.2 | 0.03<sup>a</sup> |
| L ITG | 0.11 | 0.048<sup>a</sup> |
| L MTG | 0.09 | 0.345 |
| L STG | 0.04 | 1.0 |
| R AG | 0.1 | 0.187 |
| R IPL | 0.1 | 0.32 |
| R PCU | 0.03 | 1.0 |
| R SMG | 0.1 | 0.063 |
| R SPL | -0.05 | 0.909 |
| R IOG | -0.05 | 1.0 |
| R MOG | -0.1 | 0.432 |
| R SOG | -0.08 | 0.5 |
| R calc | -0.22 | 0.03<sup>a</sup> |
| R cun | -0.08 | 0.5 |
| R fus | -0.1 | 0.364 |
| R ling | -0.18 | 0.03<sup>a</sup> |
| R ITG | 0.12 | 0.03<sup>a</sup> |
| R MTG | 0.1 | 0.063 |
| R STG | 0.13 | 0.03<sup>a</sup> |

*Reported p-values are Bonferroni-Holm corrected for multiple comparisons*
$r_s$ = Spearman's rho
<sup>a</sup> passed Bonferroni-Holm correction for multiple comparisons

**Figure 3.**
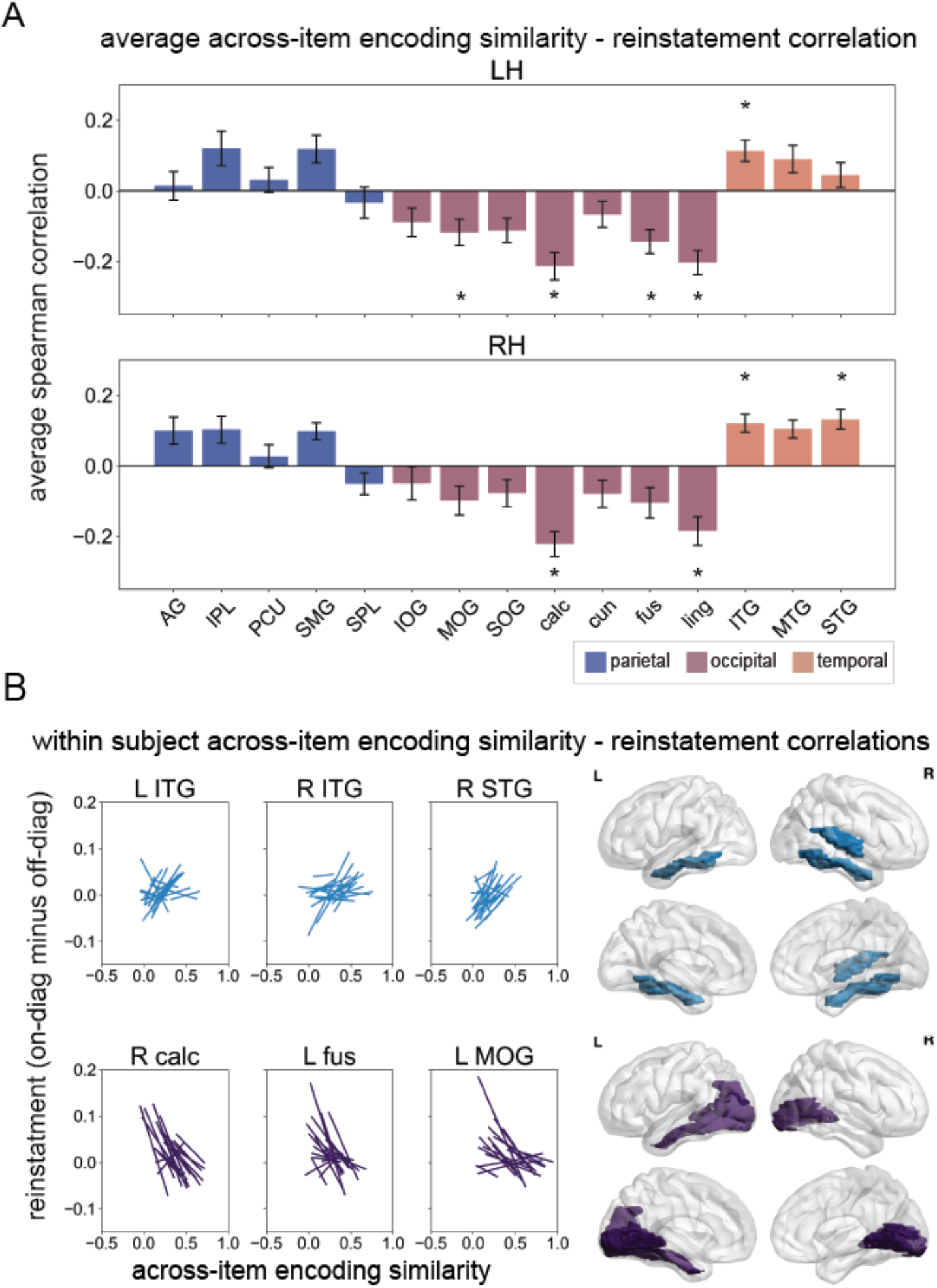
Across-item encoding similarity differentially predicts reinstatement. A) Average within-subject correlation between across-item encoding similarity and reinstatement, split by hemisphere (LH – left, RH – right). Standard error of the mean plotted as error bars. B) Within-subject correlations between across item-similarity and reinstatement. Top panel shows significant positive correlations, with each line representing an individual subject’s linear fit (reinstatement ∼ encoding similarity) for significant ROIs. Glass brains to the right show lateral (top) and medial (bottom) views of all ROIs showing a significant positive effect in blue for left (L) and right hemispheres (R). Bottom panel shows negative correlations for a subset of significant ROIs. Glass brains to the right show lateral and medial views of all ROIs showing a significant negative effect in purple.

Because the reinstatement measure reflects a difference score, it is possible that reinstatement of irrelevant videos (off-diagonal correlations) was driving the association with across-item encoding similarity rather than reinstatement of the video itself. To test for this, we repeated the same correlation analyses using reinstatement of matched pairs (diagonal) without subtracting reactivation of different videos (off-diagonal). A similar set of regions to the previous analysis showed significant effects. Across-item encoding similarity and reinstatement were positively correlated in bilateral inferior temporal gyrus, supramarginal gyrus, right middle temporal gyrus, right superior temporal gyrus, and were negatively correlated in bilateral calcarine gyrus and lingual gyrus (all p’s < 0.05, corrected for multiple comparisons).

Thus, in high-level sensory processing regions across temporal and parietal lobes, videos that are more highly integrated with other videos at encoding (more neurally average) show higher reinstatement at retrieval, while videos that are more differentiated at encoding (more neurally distinct) show lower reinstatement. Occipital regions, particularly lower-level visual processing regions (i.e. primary visual cortex), but also high-level visual processing regions (fusiform gyrus, middle occipital gyrus), show the opposite pattern, with greater integration at encoding leading to lower reinstatement and greater differentiation leading to higher reinstatement. These findings demonstrate that an event’s neural representation can be simultaneously integrated and differentiated in different populations across the brain, perhaps reflecting differences in the type of content represented across regions.

### Across-item encoding similarity is related to later memory performance

We next examined whether the degree to which videos are encoded in an integrated or differentiated fashion affects their subsequent memory performance. Memory performance was measured objectively during the retrieval stage by two true/false questions about the content of the videos, and subjectively by vividness ratings (1-4). To code for accuracy, each trial was given a score from 0-2 to represent the number of questions answered correctly. For each subject and each ROI, we computed the Kendall’s Tau rank correlation between across-item encoding similarity and each memory measure across trials.

Across-item encoding similarity in parietal and temporal regions was significantly positively correlated with accuracy at retrieval. Permutation tests revealed that this correlation was significantly greater than zero for bilateral superior parietal lobule, supramarginal gyrus, inferior temporal gyrus, superior temporal gyrus, left angular gyrus, left inferior parietal lobule, right middle temporal gyrus, and right inferior occipital gyrus ROIs (all p’s < 0.05, corrected for multiple comparisons). Average across-item encoding similarity plotted by accuracy score revealed a linear relationship, such that average neural similarity was highest for trials that were later remembered with 100% accuracy (2/2 questions), with lower similarity for trials that were remembered with 50% accuracy (1/2 questions) and 0% accuracy (0/2 questions) (Figure 4A). This indicates that videos that are highly integrated with other videos in temporal, parietal, and occipital lobes are more likely to be remembered with greater accuracy relative to videos that show low integration in these regions.

**Figure 4.**
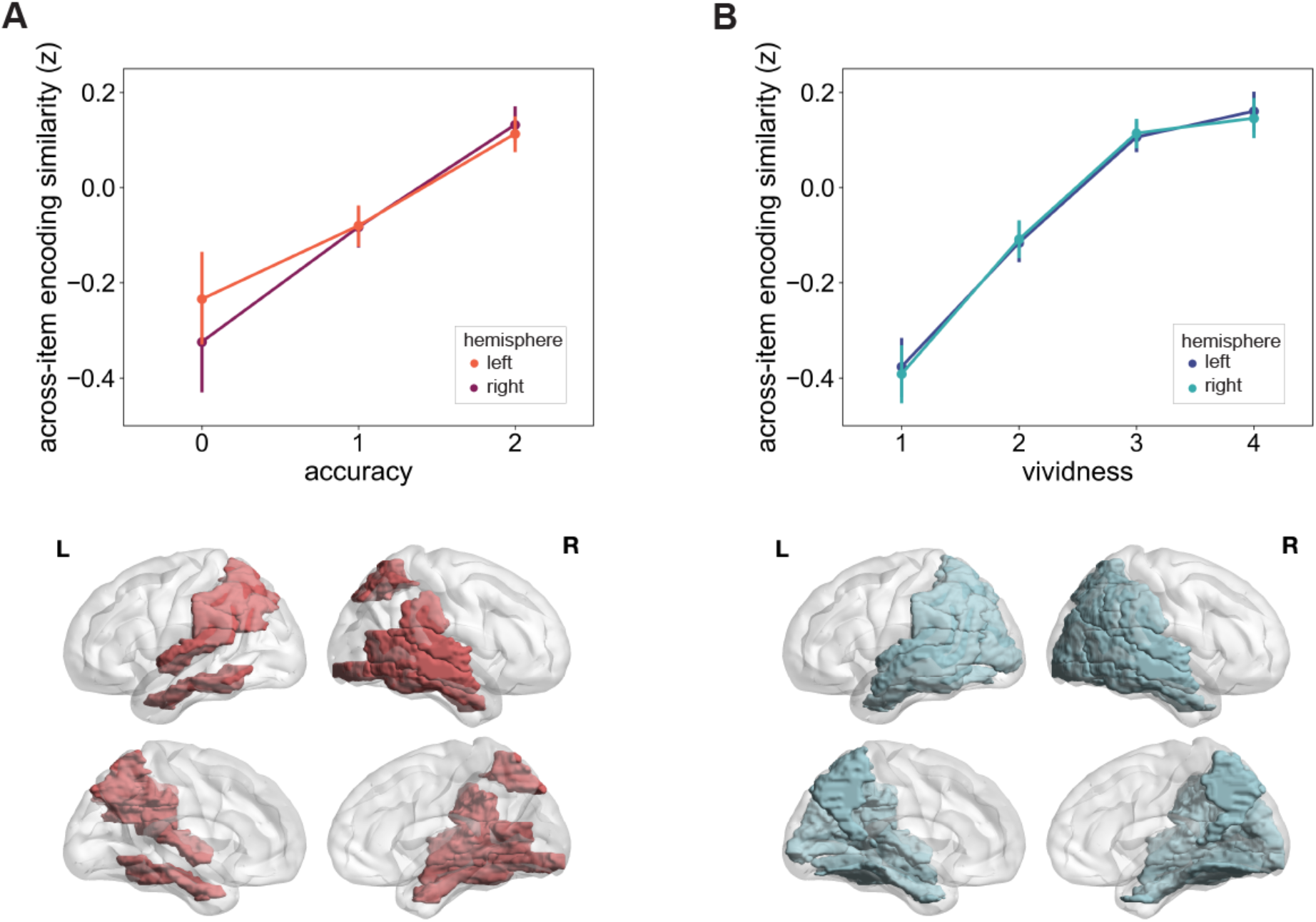
Across-item encoding similarity relates to later memory performance. A) Across-item encoding similarity plotted by accuracy score, averaged across all significant ROIs. Glass brains below show lateral (top) and medial (bottom) view of all ROIs showing a significant correlation between accuracy and across-item encoding similarity for left (L) and right hemispheres (R). B) Across-item encoding similarity plotted by vividness score, averaged across all significant ROIs. Glass brains below show lateral (top) and medial (bottom) view of all ROIs showing a significant positive effect for left and right hemispheres. Error bars reflect 95% confidence intervals. Across-item encoding similarity scores have been converted to z-scores for visualization purposes.

We next examined vividness ratings, finding that across-item encoding similarity in parietal, temporal, and occipital ROIs was positively correlated with vividness ratings at retrieval. Permutation tests revealed that the correlation between across-item encoding similarity and vividness was significantly greater than zero for bilateral angular gyrus, inferior parietal lobe, superior parietal lobe, precuneus, supramarginal gyrus, inferior temporal gyrus, middle temporal gyrus, superior temporal gyrus, middle occipital gyrus, fusiform gyrus, and right inferior and superior occipital gyri (all p’s < 0.05, corrected for multiple comparisons). Across-item encoding similarity increased with vividness ratings (Figure 4B), suggesting that videos that are more highly integrated at encoding are later remembered with more subjective vividness. Together these findings suggest that greater integration of related videos at encoding in high-level sensory processing regions across parietal, temporal, and occipital lobes is beneficial for later subjective and objective memory. Notably, low-level visual processing regions (i.e. primary visual cortex) were not significantly associated with later memory, suggesting that the degree of neural overlap between videos at encoding in these regions does not strongly predict later memory. See Table 3.

**Table 3.** Across-item encoding similarity – memory performance correlation by ROI.

| Region | Accuracy |  | Vividness |  |
| --- | --- | --- | --- | --- |
| | $\tau$ | p-value | $\tau$ | p-value |
| L AG | 0.09 | 0.044<sup>a</sup> | 0.11 | 0.03<sup>a</sup> |
| L IPL | 0.09 | 0.03<sup>a</sup> | 0.14 | 0.03<sup>a</sup> |
| L PCU | 0.09 | 0.088 | 0.16 | 0.03<sup>a</sup> |
| L SMG | 0.1 | 0.03<sup>a</sup> | 0.12 | 0.036<sup>a</sup> |
| L SPL | 0.08 | 0.044<sup>a</sup> | 0.14 | 0.03<sup>a</sup> |
| L IOG | 0.05 | 0.517 | 0.07 | 0.056 |
| L MOG | 0.05 | 0.581 | 0.11 | 0.03<sup>a</sup> |
| L SOG | 0.05 | 0.634 | 0.1 | 0.056 |
| L calc | 0.02 | 1.0 | 0.02 | 0.486 |
| L cun | -0.0 | 1.0 | 0.04 | 0.486 |
| L fus | 0.07 | 0.144 | 0.11 | 0.03<sup>a</sup> |
| L ling | 0.0 | 1.0 | 0.06 | 0.056 |
| L ITG | 0.1 | 0.044<sup>a</sup> | 0.15 | 0.03<sup>a</sup> |
| L MTG | 0.1 | 0.075 | 0.14 | 0.03<sup>a</sup> |
| L STG | 0.09 | 0.03<sup>a</sup> | 0.14 | 0.03<sup>a</sup> |
| R AG | 0.09 | 0.075 | 0.13 | 0.03<sup>a</sup> |
| R IPL | 0.07 | 0.075 | 0.14 | 0.03<sup>a</sup> |
| R PCU | 0.08 | 0.068 | 0.15 | 0.03<sup>a</sup> |
| R SMG | 0.1 | 0.044<sup>a</sup> | 0.12 | 0.03<sup>a</sup> |
| R SPL | 0.1 | 0.03<sup>a</sup> | 0.14 | 0.03<sup>a</sup> |
| R IOG | 0.08 | 0.03<sup>a</sup> | 0.1 | 0.03<sup>a</sup> |
| R MOG | 0.08 | 0.054 | 0.11 | 0.03<sup>a</sup> |
| R SOG | 0.08 | 0.075 | 0.1 | 0.03<sup>a</sup> |
| R calc | 0.06 | 0.12 | 0.04 | 0.486 |
| R cun | 0.08 | 0.068 | 0.07 | 0.12 |
| R fus | 0.05 | 0.272 | 0.1 | 0.03<sup>a</sup> |
| R ling | 0.04 | 0.703 | 0.05 | 0.232 |
| R ITG | 0.13 | 0.03<sup>a</sup> | 0.13 | 0.03<sup>a</sup> |
| R MTG | 0.11 | 0.03<sup>a</sup> | 0.13 | 0.03<sup>a</sup> |
| R STG | 0.13 | 0.03<sup>a</sup> | 0.15 | 0.03<sup>a</sup> |

*Reported p-values are Bonferroni-Holm corrected for multiple comparisons*
$\tau$ = Kendall's Tau
<sup>a</sup> *passed Bonferroni-Holm correction for multiple comparisons*

### Testing the effects of hippocampal encoding activity on later retrieval

Although the focus of the current study was on pattern similarity outside of the medial temporal lobes, previous studies have found that pattern integration or differentiation within the hippocampus predicts later memory retrieval and reinstatement (Brunec et al., 2020; LaRocque et al., 2013; Schlichting et al., 2014). While the hippocampus is thought to be important for coordinating cortical reinstatement, we did not predict that reinstatement would occur within the hippocampus itself, in line with several previous studies (Wing, 2015; Ritchey, 2013). To test whether encoding pattern similarity relates to reinstatement in the hippocampus, we used left and right hippocampus as ROIs for the same across-item pattern similarity and reinstatement analyses described above. We found no significant correlation between hippocampal across-item encoding similarity and reinstatement (left hippocampus: p = .71; right hippocampus: p = .55). We next tested whether across-item encoding similarity in the hippocampus predicts later memory, repeating the above analyses correlating encoding activity with memory performance using left and right hippocampal ROIs. There was a significant positive correlation between across-item encoding similarity in the left hippocampus and both accuracy (p = .003) and vividness ratings (p = .005), while right hippocampus was correlated with accuracy (p = .014) but not accuracy (p = .491). Thus, videos that are more highly integrated with related videos at encoding in the hippocampus are later remembered with greater accuracy and vividness, but reinstatement does not appear to occur in the hippocamps at retrieval.

### Testing the relevance of cortical reinstatement to memory performance

We next performed an exploratory analysis to see whether cortical reinstatement is related to later memory performance. For each subject, we computed the Kendall’s Tau rank correlation between reinstatement and each memory measure across trials. We then tested whether the correlation between reinstatement and memory was significantly greater than a null distribution of correlation values obtained by permutation, for each ROI. There were no significant correlations between reinstatement and either accuracy or vividness for any ROIs (all p’s > .05 after correction).

## Discussion

This study examined how highly overlapping naturalistic events are represented in cortical activity patterns at encoding and retrieval. We showed that the degree and content of visual similarity between videos affects their encoding, such that highly similar videos tend to be integrated while dissimilar videos tend to be differentiated. In high-level sensory processing regions in temporal and parietal cortices, videos that were encoded with greater overlapping patterns of activity, suggesting integration, showed higher reinstatement at retrieval. Greater integration in high-level sensory processing regions also predicted later memory performance. Conversely, in visual processing regions in occipital cortex, videos that were encoded in more distinctive patterns of neural activity, suggesting differentiation, showed greater reinstatement. These findings demonstrate that events can be simultaneously integrated and differentiated in different neural populations, with effects on later memory differing by brain region.

Current evidence suggests that similarity between events may influence whether they are integrated or differentiated at encoding (Cox et al., 2021; Koen & Rugg, 2016). We found that events with high visual similarity to other events tended to be encoded in overlapping patterns of neural activity in parietal, occipital, and temporal cortices, suggesting that visually similar videos are integrated in sensory processing regions throughout the cortex. Similarity between low-level visual features predicted integration in high- and low-level sensory processing regions predominantly in occipital cortex, and similarity between higher-level visual features predicted integration in a high-level visual processing region in temporal cortex. Visual features from CNNs have been shown to predict neural representations in the visual cortex, with early and late CNN layers mapping onto early and late visual cortex, respectively (Güçlü & van Gerven, 2015; Yamins et al., 2014). One possible explanation for our finding that early layer features predicted representations in both early and late visual cortex is that, due to the hierarchical organization of the visual cortex, representations in later visual cortex regions are based off early visual cortex representations (Serre, 2014). Representations in higher-level sensory processing regions may therefore reflect more complex visual features in addition to the basic features from which they are derived. Overall, these findings indicate that the content of visual overlap between events affects their encoding, with highly similar events tending to be integrated throughout cortical regions.

Previous studies have shown that neural activity when encoding a stimulus is related to the strength of its cortical reinstatement at recall (Gordon et al., 2014; Hebscher et al., 2021; Ritchey et al., 2013; Wing et al., 2015). We found that neural similarity between encoded events differentially predicted their later reinstatement across the cortex. Greater similarity between videos at encoding predicted stronger reinstatement in lateral temporal and parietal regions. Lateral temporal regions have been implicated in semantic processing and memory, action perception, and high-level visual processing such as object, face, and scene perception (Conway, 2018; Kable et al., 2005; Rogers et al., 2006). Lateral parietal cortex plays a role in multisensory integration, schema integration, and representing stimulus-specific details important for subjective memory vividness (Kuhl & Chun, 2014; Lee & Kuhl, 2016; Pasalar et al., 2010; Wagner et al., 2015). Our findings suggest that integrating the high-level sensory and semantic content of videos is beneficial for later reinstatement of this content. Future studies could directly test whether similarity between high-level features of events affects their encoding and reinstatement by manipulating stimuli to explicitly differ on these features.

Occipital cortex regions showed the opposite effect, with greater neural similarity between videos at encoding predicting lower reinstatement, or greater dissimilarity between events leading to stronger reinstatement. This effect was found in calcarine and lingual gyri, which are implicated in processing of basic visual features, as well as left fusiform gyrus and middle occipital gyrus, which play roles in higher-level visual functions such as mental imagery, and face and object perception (Spagna et al., 2021; Tootell et al., 1998; Tu et al., 2013). Many of these regions also showed a positive relationship between low-level visual similarity and neural similarity at encoding, indicating that videos with high overlap between basic visual features tend to encoded in overlapping patterns of neural activity at encoding, which then leads to lower reinstatement in occipital cortex. Interestingly, the nature of this negative association was such that greater integration in occipital cortex tended to be associated with more negative reinstatement. Some previous studies have reported a negative correlation between univariate activity at encoding and retrieval in posteromedial cortex, reflecting beneficial deactivation at encoding and activation at retrieval (Gilmore et al., 2015; Vannini et al., 2011). It is therefore possible that a negative encoding-retrieval correlation in occipital cortex (i.e. negative reinstatement) still reflects reinstatement of visual content, with visual representations being activated during encoding (i.e. perception) and deactivated at retrieval (i.e. mental imagery). Indeed, visual cortex has been shown to deactivate during memory and mental imagery (Mazard et al., 2004). However, the functional relevance of a negative encoding-retrieval similarity correlation in the present study remains unclear. Additional work is needed to clarify the nature of the negative association between across-item encoding similarity and reinstatement in occipital regions.

Previous studies have found that both cortical integration and differentiation is beneficial for later episodic memory performance (Chanales et al., 2019; Katsumi et al., 2021; Koen & Rugg, 2016; Wing et al., 2020). Here we showed that accuracy and vividness of retrieved memories are predicted by integration across different cortical regions. Videos that were encoded with overlapping neural representations in lateral parietal cortex, temporal cortex, and right occipital gyri were more likely to be remembered with greater accuracy. These regions are broadly associated with semantic processing, high-level visual processing, multimodal integration, and forming stimulus-specific representations for memory retrieval (Bonnici et al., 2016; Conway, 2018; Lee & Kuhl, 2016; Rogers et al., 2006; Tu et al., 2013). Our findings therefore suggest that integrating high-level features in lateral temporal and parietal cortex, as well as occipital cortex, allows participants to recall the precise details of events with greater objective accuracy. Subjective vividness of memories was predicted by a wider set of cortical regions encompassing medial and lateral parietal lobe, temporal lobe, and occipital lobe. Most of the same regions that showed an association with accuracy also predicted vividness, in addition to several other regions such as precuneus, fusiform gyrus, and superior occipital gyri. In line with this finding, episodic memory vividness has been associated with neural activity in a wide network of regions, including precuneus, angular gyrus, and occipitotemporal regions (Geib et al., 2017; Hebscher et al., 2019; Richter et al., 2016; Tibon et al., 2019). Our results show that forming overlapping neural representations of similar naturalistic events in the cortex is beneficial for later memory.

We also found that reinstatement does not predict memory, which calls into question the notion that reinstatement reflects memory retrieval (Bainbridge et al., 2021; Hebscher et al., 2021; Rissman & Wagner, 2012). Reinstatement within cortical regions is thought to reflect the retrieval of region-specific content of an event (Danker & Anderson, 2010). Reinstatement within an isolated cortical region may therefore not map onto overt memory performance, which is likely based on more global representations of events. Future studies could investigate how reinstatement across a network of cortical regions influences memory performance above and beyond reinstatement in specific regions.

Taken together, our findings indicate regional dissociations in how highly similar naturalistic stimuli are encoded and retrieved. In parietal and temporal lobes, integrating similar stimuli at encoding leads to stronger reinstatement and better objective and subjective memory for those stimuli. In high-level visual processing regions in occipital cortex, integration leads to lower or more negative reinstatement and is predictive of later subjective memory. Conversely, in low-level visual cortex regions, integration predicts lower reinstatement not memory performance. These findings suggest that integrating high-level features at encoding leads to better memory but differential reinstatement of those features throughout the cortex. Integrating low-level visual features leads to lower reinstatement of those features in early visual cortex but does not affect memory, possibly because memory in this task is complex and depends on multiple higher-order functions. These divergent effects may reflect the brain’s ability to simultaneously integrate high-level features of events and distinguish overlapping low-level features, which can allow us to remember both relationships between events and their specific details (Dimsdale-Zucker et al., 2018). In conclusion, our findings show that encoding-related integration and differentiation processes across the cortex have divergent effects on later recall of highly similar naturalistic events.

## Methods

### Participants

Twenty adults participated in the current study (11 females, mean age = 24.26, SD = 3.57, range = 19-32). Data from these same participants has been published elsewhere, in a study that examined the effects of transcranial magnetic stimulation (TMS) on hippocampal coordination of cortical reinstatement (Hebscher et al., 2021). All participants were native or fluent English speakers, had normal or corrected-to-normal vision, and were free from a history of neurological illness or injury, psychiatric condition, substance abuse, or serious medical conditions. All participants passed standard MRI safety screenings. Participants provided informed consent prior to participating in the experiment and were paid for participation. Study procedures were approved by the Northwestern University Institutional Review Board.

### Experimental design

The data reported in the present study were originally collected as a control session for a previously published TMS study (Hebscher et al., 2021). During control sessions, participants received continuous theta burst TMS to their vertex for 60 s, approximately 7 minutes prior to beginning an episodic memory task while fMRI was collected. Vertex stimulation is believed to have no effect on episodic memory or related brain activity.

### Episodic memory task

Participants performed a practice version of the episodic memory task with unique stimuli prior to entering the MRI scanner to familiarize them with the task and to ensure correct performance. Once moved to the MRI scanner, participants completed the episodic memory task in which they watched and recalled a series of short video clips (Figure 1A). Videos consisted of 51 short (7 s) depictions of common events such as draining pasta, kicking a ball, and putting a sheet on the bed. Each video depicted a unique event centered around the same one character and occurred in a fixed number of locations, resulting in high overlap between elements of the videos. All videos were presented without sound. Videos were presented in a randomized order at both encoding and retrieval phases.

At the start of the encoding phase, participants were reminded of the task instructions and were instructed to pay close attention to all elements of the videos as their memory for the videos would later be tested. Each video was then presented alone for 7 s, followed by an interstimulus interval (ISI) of 1 s. Participants then judged whether a number was odd or even for 2 s to discourage continued processing of the video. Previous evidence suggests that such judgments drive activity in key regions such as hippocampus back to zero during the intertrial interval (ITI) (Stark & Squire, 2001). The trial ended with an ITI (fixation cross), jittered at 4, 6, and 8 s. Each encoding trial was 16 s on average. All encoding trials occurred within 1 fMRI run lasting 13.9 mins.

The retrieval phase began immediately following completion of the encoding phase. Following a reminder of the instructions for this phase, participants were presented with the description of a video for 3 s and told to mentally replay the video within the allotted time, within a blank box that remained on the screen alone for 7 s. After mentally recalling each video, participants were given 4s to rate their vividness of the memory on a scale from 1-4, with 1 meaning they did not recall the video at all, and 4 meaning they recalled it vividly. Following the vividness rating, participants answered two true/false (yes/no) accuracy questions about the videos, incorporating their confidence (1-definitely yes, 2-maybe yes, 3-maybe no, 4-definitely no). 5 s were allotted for each accuracy question. There was a total of 1.2 s ISI within each trial, and an ITI (fixation cross) jittered at 4, 6, and 8 s, with an average trial length of 34 s. Responses were made using 2 response boxes, each with 2 buttons. Retrieval trials occurred over 3 fMRI runs (17 trials each), each lasting 10 mins. Average vividness scores were 2.71 (SD = 0.36), and accuracy was on average 70% (question 1 = 0.73 (0.09); question 2 = 0.67 (0.07)). Means and standard deviations on all episodic memory measures are reported in a previous study (Hebscher et al., 2021).

### MRI acquisition

Structural and functional images were acquired using a Siemens 3 T Prisma whole-body scanner with a 64-channel head coil located in the Northwestern University Center for Translational Imaging Facility. Functional images were acquired using a whole-brain BOLD EPI sequence (TR = 2000 ms, TE = 20 ms, FOV = 1116×1080 mm, flip angle = 80°, and 1.7×1.7×1.7 mm voxel resolution, over 275 volumes). Structural images were acquired using a T1-weighted MPRAGE sequence (TR = 2170 ms, TE = 1.69 ms, FOV = 256×256 mm, flip angle = 7°, voxel resolution: 1.0 × 1.0 × 1.0 mm, 1-mm thick sagittal slices). The encoding phase of the memory task consisted of one 13.9 min run (418 volumes), while retrieval was split among three 10 min runs (299 volumes each).

### MRI preprocessing

Functional MRI data were preprocessed using AFNI software. Preprocessing included functional-structural co-registration, motion correction, spatial smoothing using a 1.7-mm full-width-half-maximum (FWHM) isotropic Gaussian kernel, and signal intensity normalization by the mean of each voxel. Motion parameters were calculated for each volume, and volumes with excessive frame to frame displacement (>0.3 mm) were flagged for later censoring.

Single-trial estimates were generated for multivariate analyses using a general linear model (GLM) in AFNI (3dDeconvolve). A separate model was constructed for each individual trial to estimate its activity separately from all other trials and nuisance variables, an approach known to work effectively for single-trial estimation for multivoxel pattern analyses (Mumford et al, 2012). For each functional run, individual trials were modelled separately against all other trials using a response model of a 7-s block convolved with a canonical hemodynamic response function. Nuisance variables included the six affine motion estimates generated by motion correction as well as linear drift. For encoding trials, the 7-s block began at the start of video presentation, while at retrieval the 7-s block began when participants saw the cued video title. The resulting single-trial t-maps for each trial were used for subsequent analyses, based on recommendations that using t-maps rather than beta maps for representational similarity analyses reduces the influence of noisy voxels (Dimsdale-Zucker & Ranganath, 2019). All multivariate analyses were carried out in native space.

### Regions of interest

Multivariate analyses were focused on a set of 15 bilateral ROIs spanning parietal, temporal, and occipital cortices (Figure 1C). Cortical ROIs included superior parietal lobule, inferior parietal lobule, supramarginal gyrus, angular gyrus, precuneus, calcarine gyrus, cuneus, lingual gyrus, superior occipital gyrus, middle occipital gyrus, inferior occipital gyrus, fusiform gyrus, superior temporal gyrus, middle temporal gyrus, inferior temporal gyrus. A hippocampal ROI was also used for control analyses. ROIs were defined using the Eickhoff-Zilles macro labels from N27 in MNI space (CA_ML_18_MNI atlas in AFNI) on a template brain in MNI space and warped to native space for each participant by calculating the transformation matrix needed to warp into MNI space and then reverse-transforming all ROIs into native space using affine transformation (3dAllineate).

### Representational similarity analysis

#### Across-item encoding pattern similarity

We examined similarity between neural representations of different videos at encoding by conducting a series of representational similarity analyses (RSAs) on patterns of neural activity within ROIs. RSAs measured the Pearson’s correlation of each video’s evoked fMRI activity at encoding with the activity pattern of every other video at encoding, for each ROI. Pairwise correlations between all videos resulted in an encoding-encoding similarity matrix, with the diagonal reflecting each video’s correlation with itself (1.0), and the off-diagonal reflecting each video’s correlation with every other video. Pairwise correlation values were then Fisher transformed, and for each video (row), we calculated the average correlation with all other videos. This measure, termed across-item encoding similarity, reflects the relative similarity of a given video to all other videos, effectively its place on a continuum from neural averageness to distinctness.

#### Encoding-retrieval reinstatement

We examined reinstatement of video-specific patterns of neural activity by conducting a series of RSAs on patterns of neural activity within ROIs. RSAs measured encoding-retrieval similarity by computing the correlation between neural activity patterns for all pairs of videos, for each ROI. Only trials that were self-reported as recollected (vividness score of > 1) were included in these analyses (86% trials). Pairwise correlations between all videos resulted in a matrix with the diagonal reflecting correlations between the same video at encoding and retrieval (matched pairs), and the off-diagonal reflecting correlations between different videos (mismatched pairs). Pairwise correlation values were Fisher transformed. We then subtracted the mean of the off-diagonal correlation values from the mean of the diagonal correlations and took the average of these on-vs. off-diagonal correlation differences. This metric reflects the degree to which neural pattern similarity was greater for matched versus mismatched video pairs.

RSAs for both encoding-encoding and encoding-retrieval similarity were computed in MATLAB (The MathWorks, Inc., Natick, MA, USA), using the CoSMoMVPA toolbox (cosmomvpa.org) (Oosterhof et al., 2016) functions to load and organize fMRI datasets (cosmo_fmri_dataset), calculate encoding-encoding similarity matrices (cosmo_dissimilarity_matrix_measure), and encoding-retrieval similarity matrices (cosmo_correlation_measure). All subsequent analyses on correlation matrices were performed using in-house MATLAB scripts.

### Convolutional neural network measure of visual similarity

To obtain a measure of visual similarity, we extracted learned image features from the pretrained CNN AlexNet (Krizhevsky et al., 2012) (Figure 1D). CNNs are deep learning algorithms inspired by the organization of the visual cortex that consist of multiple layers, where later layers represent increasingly complex stimulus features (Zeiler & Fergus, 2014). AlexNet is a CNN that is 8 layers deep and is trained on more than 1 million images to classify images into 1000 object categories. Briefly, AlexNet consists of 5 convolutional layers and 3 fully connected layers. Each convolutional layer integrates inputs from the immediate previous later and encodes them in more compact activity patterns. The size of filter (kernel) shrinks with convolutional layers, while the number of filters increases and their receptive field enlarges, allowing deeper layers to take in larger-scale, more comprehensive features. In fully connected layers, each neuron is connected to all neurons in previous layers with its own weights. For a more complete description of AlexNet’s construction, see Krizhevsky et al. (2012). One frame from the midpoint of each video was used as input for the pretrained network. We then extracted features from two layers of the network, an early layer (convolutional layer 2; conv2), reflecting a lower-level representation of each image, and a deeper layer (fully connected layer 7; fc7), reflecting higher-level features constructed using the lower-level features of earlier layers. Early and deep layers are thought to correspond to visual information from early and late visual cortex, respectively (Güçlü & van Gerven, 2015; Yamins et al., 2014). The early layer (conv2) had a total of 186,624 features, while the deep layer (fc7) had 4,096 features. For each layer, we constructed a correlation matrix of the Pearson correlation between each video’s features. Visual similarity for each video was computed by taking the average Fisher-transformed off-diagonal correlation for each video, or the average of each video’s correlation with all other videos. AlexNet was implemented in MATLAB using the Deep Learning Toolbox.

### Group-level analyses

Group-level statistics were performed to assess the relationship between (1) across-item encoding similarity and visual similarity, (2) across-item encoding similarity and reinstatement, and (3) across-item encoding similarity and memory performance. For each participant and ROI, we measured the correlation between the two measures of interest. One-sample permutation tests were then used to assess the significance of the correlation values at the group-level. These tests create a null distribution by assigning random signs to the observed correlation values and recomputing the difference in means from the null population mean 1000 times. Resulting p–values reflect the probability of the absolute value of observed correlation differences being more extreme than the absolute value of the permuted differences. Familywise error rates were corrected for multiple comparisons using Bonferroni-Holm method (Holm, 1979). Corrected p-values are reported.

## References

Bainbridge, W. A., Hall, E. H., & Baker, C. I. (2021). Distinct Representational Structure and Localization for Visual Encoding and Recall during Visual Imagery. Cerebral Cortex, 31(4), 1898–1913. https://doi.org/10.1093/cercor/bhaa329

Bonnici, H. M., Richter, F. R., Yazar, Y., & Simons, J. S. (2016). Multimodal Feature Integration in the Angular Gyrus during Episodic and Semantic Retrieval. Journal of Neuroscience, 36(20), 5462–5471. https://doi.org/10.1523/JNEUROSCI.4310-15.2016

Brunec, I. K., Robin, J., Olsen, R. K., Moscovitch, M., & Barense, M. D. (2020). Integration and differentiation of hippocampal memory traces. Neuroscience and Biobehavioral Reviews, 118, 196–208. https://doi.org/10.1016/j.phrs.2020.104743

Chanales, A. J. H., Dudukovic, N. M., Richter, F. R., & Kuhl, B. A. (2019). Interference between overlapping memories is predicted by neural states during learning. Nature Communications, 10(1), 5363. https://doi.org/10.1038/s41467-019-13377-x

Conway, B. R. (2018). The organization and operation of inferior temporal cortex. Annual Review of Vision Science, 4, 381.

Cox, W. R., Dobbelaar, S., Meeter, M., Kindt, M., & van Ast, V. A. (2021). Episodic memory enhancement versus impairment is determined by contextual similarity across events. Proceedings of the National Academy of Sciences, 118(48). https://doi.org/10.1073/pnas.2101509118/-/DCSupplemental

Danker, J. F., & Anderson, J. R. (2010). The ghosts of brain states past: remembering reactivates the brain regions engaged during encoding. Psychological Bulletin, 136(1), 87–102. https://doi.org/10.1037/a0017937

Dimsdale-Zucker, H. R., & Ranganath, C. (2019). Representational Similarity Analyses: A Practical Guide for Functional MRI Applications. Handbook of Behavioral Neuroscience, 28, 509–525. https://doi.org/10.1016/B978-0-12-812028-6.00027-6

Dimsdale-Zucker, H. R., Ritchey, M., Ekstrom, A. D., Yonelinas, A. P., & Ranganath, C. (2018). CA1 and CA3 differentially support spontaneous retrieval of episodic contexts within human hippocampal subfields. Nature Communications, 9(1), 294. https://doi.org/10.1038/s41467-017-02752-1

Ezzyat, Y., Inhoff, M. C., & Davachi, L. (2018). Differentiation of human medial prefrontal cortex activity underlies long-term resistance to forgetting in memory. Journal of Neuroscience, 38(48), 10244–10254.

Favila, S. E., Chanales, A. J. H., & Kuhl, B. A. (2016). Experience-dependent hippocampal pattern differentiation prevents interference during subsequent learning. Nature Communications, 7. https://doi.org/10.1038/ncomms11066

Geib, B. R., Stanley, M. L., Wing, E. A., Laurienti, P. J., & Cabeza, R. (2017). Hippocampal contributions to the large-scale episodic memory network predict vivid visual memories. Cerebral Cortex, 27(1), 680–693.

Gilmore, A. W., Nelson, S. M., & McDermott, K. B. (2015). A parietal memory network revealed by multiple MRI methods. Trends in Cognitive Sciences. https://doi.org/10.1016/j.tics.2015.07.004

Gordon, A. M., Rissman, J., Kiani, R., & Wagner, A. D. (2014). Cortical reinstatement mediates the relationship between content-specific encoding activity and subsequent recollection decisions. Cerebral Cortex, 24(12), 3350–3364. https://doi.org/10.1093/cercor/bht194

Güçlü, U., & van Gerven, M. A. J. (2015). Deep Neural Networks Reveal a Gradient in the Complexity of Neural Representations across the Ventral Stream. https://doi.org/10.6080/K0QN64NG

Hebscher, M., Kragel, J. E., Kahnt, T., & Voss, J. L. (2021). Enhanced reinstatement of naturalistic event memories due to hippocampal-network-targeted stimulation. Current Biology, 31(7), 1428–1437. https://doi.org/10.1101/2020.08.18.256008

Hebscher, M., Meltzer, J., & Gilboa, A. (2019). A causal role for the precuneus in network-wide theta and gamma oscillatory activity during complex memory retrieval. Elife, 8, e43114.

Holm, S. (1979). A simple sequentially rejective multiple test procedure. Scandinavian Journal of Statistics, 65–70.

Katsumi, Y., Andreano, J. M., Barrett, L. F., Dickerson, B. C., & Touroutoglou, A. (2021). Greater Neural Differentiation in the Ventral Visual Cortex Is Associated with Youthful Memory in Superaging. 1–13.

Koen, J. D., & Rugg, M. D. (2016). Memory reactivation predicts resistance to retroactive interference: Evidence from multivariate classification and pattern similarity analyses. Journal of Neuroscience, 36(15), 4389–4399. https://doi.org/10.1523/JNEUROSCI.4099-15.2016

Kriegeskorte, N., Mur, M., & Bandettini, P. A. (2008). Representational similarity analysis-connecting the branches of systems neuroscience. Frontiers in Systems Neuroscience, 2, 4.

Krizhevsky, A., Sutskever, I., & Hinton, G. E. (2012). Imagenet classification with deep convolutional neural networks. Advances in Neural Information Processing Systems, 1097–1105.

Kuhl, B. A., & Chun, M. M. (2014). Successful remembering elicits event-specific activity patterns in lateral parietal cortex. Journal of Neuroscience, 34(23), 8051–8060.

LaRocque, K. F., Smith, M. E., Carr, V. A., Witthoft, N., Grill-Spector, K., & Wagner, A. D. (2013). Global similarity and pattern separation in the human medial temporal lobe predict subsequent memory. Journal of Neuroscience, 33(13), 5466–5474. https://doi.org/10.1523/JNEUROSCI.4293-12.2013

Lee, H., & Kuhl, B. A. (2016). Reconstructing perceived and retrieved faces from activity patterns in lateral parietal cortex. Journal of Neuroscience, 36(22), 6069–6082.

Mazard, A., Tzourio-Mazoyer, N., Crivello, F., Mazoyer, B., & Mellet, E. (2004). A PET meta-analysis of object and spatial mental imagery. European Journal of Cognitive Psychology, 16(5), 673–695.

Oedekoven, C. S. H., Keidel, J. L., Berens, S. C., & Bird, C. M. (2017). Reinstatement of memory representations for lifelike events over the course of a week. Scientific Reports, June, 1–12. https://doi.org/10.1038/s41598-017-13938-4

Oosterhof, N. N., Connolly, A. C., & Haxby, J. V. (2016). CoSMoMVPA: multi-modal multivariate pattern analysis of neuroimaging data in Matlab/GNU Octave. Frontiers in Neuroinformatics, 10, 27.

Pasalar, S., Ro, T., & Beauchamp, M. S. (2010). TMS of posterior parietal cortex disrupts visual tactile multisensory integration. European Journal of Neuroscience, 31(10), 1783–1790.

Richter, F. R., Cooper, R. A., Bays, P. M., & Simons, J. S. (2016). Distinct neural mechanisms underlie the success, precision, and vividness of episodic memory. ELife, 5, 1–18. https://doi.org/10.7554/eLife.18260

Rissman, J., & Wagner, A. D. (2012). Distributed Representations in Memory: Insights from Functional Brain Imaging. Annual Review of Psychology, 63(1), 101–128. https://doi.org/10.1146/annurev-psych-120710-100344

Ritchey, M., Wing, E. A., LaBar, K. S., & Cabeza, R. (2013). Neural similarity between encoding and retrieval is related to memory via hippocampal interactions. Cerebral Cortex (New York, N.Y. : 1991), 23(12), 2818–2828. https://doi.org/10.1093/cercor/bhs258

Rogers, T. T., Hocking, J., Noppeney, U. T. A., Mechelli, A., Gorno-Tempini, M. L., Patterson, K., & Price, C. J. (2006). Anterior temporal cortex and semantic memory: reconciling findings from neuropsychology and functional imaging. Cognitive, Affective, & Behavioral Neuroscience, 6(3), 201–213.

Schlichting, M. L., Zeithamova, D., & Preston, A. R. (2014). CA1 subfield contributions to memory integration and inference. Hippocampus, 24(10), 1248–1260.

Serre, T. (2014). Hierarchical Models of the Visual System. Encyclopedia of Computational Neuroscience, 6, 1–12.

Silson, E. H., Steel, A., Kidder, A., Gilmore, A. W., & Baker, C. I. (2019). Distinct subdivisions of human medial parietal cortex support recollection of people and places. Elife, 8.

Spagna, A., Hajhajate, D., Liu, J., & Bartolomeo, P. (2021). Visual mental imagery engages the left fusiform gyrus, but not the early visual cortex: A meta-analysis of neuroimaging evidence. Neuroscience & Biobehavioral Reviews, 122, 201–217.

Stark, C. E. L., & Squire, L. R. (2001). When zero is not zero: the problem of ambiguous baseline conditions in fMRI. Proceedings of the National Academy of Sciences, 98(22), 12760–12766.

Tibon, R., Fuhrmann, D., Levy, D. A., Simons, J. S., & Henson, R. N. (2019). Multimodal integration and vividness in the angular gyrus during episodic encoding and retrieval. Journal of Neuroscience, 39(22), 4365–4374. https://doi.org/10.1523/JNEUROSCI.2102-18.2018

Tootell, R. B. H., Hadjikhani, N. K., Mendola, J. D., Marrett, S., & Dale, A. M. (1998). From retinotopy to recognition: fMRI in human visual cortex. Trends in Cognitive Sciences, 2(5), 174–183.

Tu, S., Qiu, J., Martens, U., & Zhang, Q. (2013). Category-selective attention modulates unconscious processes in the middle occipital gyrus. Consciousness and Cognition, 22(2), 479–485.

Vannini, P., O’Brien, J., O’Keefe, K., Pihlajamäki, M., Laviolette, P., & Sperling, R. A. (2011). What goes down must come up: role of the posteromedial cortices in encoding and retrieval. Cerebral Cortex, 21(1), 22–34.

Wagner, I. C., van Buuren, M., Kroes, M. C. W., Gutteling, T. P., van der Linden, M., Morris, R. G., & Fernández, G. (2015). Schematic memory components converge within angular gyrus during retrieval. Elife, 4, e09668.

Wing, E. A., Geib, B. R., Wang, W. C., Monge, Z., Davis, X. S. W., & Cabeza, R. (2020). Cortical overlap and cortical-hippocampal interactions predict subsequent true and false memory. Journal of Neuroscience, 40(9), 1920–1930. https://doi.org/10.1523/JNEUROSCI.1766-19.2020

Wing, E. A., Ritchey, M., & Cabeza, R. (2015). Reinstatement of Individual Past Events Revealed by the Similarity of Distributed Activation Patterns during Encoding and Retrieval. Journal of Cognitive Neuroscience, 27(4), 679–691. https://doi.org/10.1162/jocn

Xue, G. (2018). The neural representations underlying human episodic memory. Trends in Cognitive Sciences, 22(6), 544–561.

Yamins, D. L. K., Hong, H., Cadieu, C. F., Solomon, E. A., Seibert, D., & DiCarlo, J. J. (2014). Performance-optimized hierarchical models predict neural responses in higher visual cortex. Proceedings of the National Academy of Sciences of the United States of America, 111(23), 8619–8624. https://doi.org/10.1073/pnas.1403112111

Zeiler, M. D., & Fergus, R. (2014). Visualizing and Understanding Convolutional Networks. European Conference on Computer Vision.

